# Expanding EMC Foldopathies: Topogenesis Deficits Alter the Neural Crest

**DOI:** 10.1101/2022.05.16.492013

**Authors:** Jonathan Marquez, Mustafa K. Khokha

## Abstract

The endoplasmic reticulum membrane protein complex (EMC) is essential for the insertion of a wide variety of transmembrane proteins into the plasma membrane across cell types. Each EMC is composed of Emc1-7, Emc10, and either Emc8 or Emc9. Recent human genetics studies have implicated variants in EMC genes as the basis for a group of human congenital diseases. The patient phenotypes are varied but appear to affect a subset of tissues more prominently than others. Namely, craniofacial development seems to be commonly affected. We previously developed an array of assays in Xenopus tropicalis to assess the effects of emc1 depletion on the neural crest, craniofacial cartilage, and neuromuscular function. We sought to extend this approach to additional EMC components identified in patients with congenital malformations. Through this approach we determine that EMC9 and EMC10 are important for neural crest development and the development of craniofacial structures. The phenotypes observed in patients and our Xenopus model were similar to EMC1 loss of function likely due to a similar mechanism of dysfunction in transmembrane protein topogenesis.

## Materials and Methods

### Xenopus Husbandry

Xenopus tropicalis were housed and cared for according to established protocols approved by Yale IACUC. We induced ovulation and collected embryos by in vitro fertilization as previously described (Lane and Khokha 2021). Embryos were raised in 1/9xMR + gentamycin. Staging of Xenopus tadpoles was performed as previously catalogued (Nieuwkoop et al. 1967).

### Xenopus CRISPR Manipulations

CRISPR/Cas9-mediated genome editing in Xenopus tropicalis embryos was used as previously described (Bhattacharya et al. 2015). CRISPR sgRNAs targeting exon 2 of emc9 and exon 1 of emc10 were designed to generate F0 knockout embryos (emc9: 5’-GGATTGGGATACAGTCACTC-3’, emc10: 5’-GGGCTGCCGGTTGTTTAGTT-3’). For targeted loss of function experiments 200 pg sgRNA along with 0.8 ng Cas9 (CP03, PNA Bio) in a 1 nl volume were injected into one cell of a two-cell stage embryo or 400 pg sgRNA along with 1.6 ng Cas9 (CP03, PNA Bio) in a 2 nl volume were injected into a one-cell stage embryo.

### Whole Mount in situ Hybridization

Whole-mount in situ Hybridization (WISH) was carried out as previously described (Henriquez et al. 1995). Briefly Xenopus embryos were fixed in 4% paraformaldehyde and dehydrated through washes in methanol. Embryos were then rehydrated in PBS with 0.1% tween-20. Embryos were then hybridized with digoxigenin labeled RNA probes complementary to target genes. Embryos were then washed and blocked prior to incubation with anti-DIG-Fab fragments (Roche) overnight at 4 degrees. BM purple (Sigma) was used to visualize expression prior to post-fixation in 4% paraformaldehyde with 0.1% glutaraldehyde.

### Whole Mount Alcian Blue Cartilage Staining

Stage 45 embryos were fixed in 100% ethanol for 48hrs at room temperature and then washed briefly in acid alcohol (1.2% HCl in 70% EtOH). A 0.25% Alcian blue solution in acid alcohol was used to stain the embryos over 48hrs at room temperature. Specimens were then washed in acid alcohol several times, rehydrated in water and bleached for 2 hours in 1.2% hydrogen peroxide under a bright (2500 lux) light. They were then washed several times in 2% KOH and left rocking overnight in 10% glycerol in 2% KOH. Samples were processed through 20%, 40%, 60 and 80% glycerol in 2% KOH.

### Motility Assay

Motility was assessed as previously described (Marquez et al. 2020). Briefly, stage 45 tadpoles were placed into separate wells of a 48-well culture dish and allowed to reach a resting state for 5 minutes. Tadpoles were then gently prodded at the rostral most aspect of the tail using a capillary pipette tip. Video of tadpole movement was captured over 30 seconds after this stimulation using an AccuScope Excelis camera mounted on a Nikon SMZ 745T stereomicroscope. Videos were analyzed using Kinovea 0.8.15 software (https://www.kinovea.org/). Analysis consisted of marking a center point of the tadpole head in each frame of a 30-second video and plotting this on a circular map. Average motion was determined via converting marked tracking points to vectors and measuring the sum of these vectors over the 30-second recorded time.

### Immunofluorescence and Microscopy

Xenopus tails were first fixed in 4% PFA for 30 minutes followed by brief permeabilization in 0.1% tween-20 in PBS. Samples were mounted in Prolong Glass (Thermo Fisher Scientific). Immunostained tails were imaged on a Zeiss Observer outfitted with optical interference (Apotome) microscopy. Antibodies are listed in Supplemental Table 1. Fluorescent images were processed and analyzed with FIJI. nAChR signals were compared via selecting 100μm x 50μm regions encompassing neuromuscular bands in the proximal Xenopus tail samples imaged. Whole-mounted, craniofacial cartilage, WISH, and TUNEL stained embryos were imaged with a Canon EOS 5d digital camera mounted on a Zeiss discovery V8 stereomicroscope.

### Immunoblot Analysis

Pooled embryos were lysed in RIPA buffer and immunoblots were performed with Bolt 4%–12% Bis-Tris plus gels and running buffer (Thermo Fisher Scientific) using standard methods. Antibodies are listed in Supplemental Table 1. Nuclear fractionation was performed using centrifugation at 720 xg for 5 min and collecting the nuclear pellet which was then washed with 500 μL of fractionation buffer (HEPES (pH 7.4) 20mM, KCl 10mM, MgCl2 2mM, EDTA 1mM, EGTA 1mM, DTT 1mM). The pellet was then dispersed via pipetting and centrifuged again at 720 xg for 10 min discarding the supernatant and resuspending the pellet in TBS with 0.1% SDS.

## Results

Our previous work evaluated the effects of emc1 loss of function on development and uncovered significant deficits in transmembrane protein topogenesis and subsequent NCC function (Marquez et al. 2020). Because of the critical importance of the EMC in inserting transmembrane proteins, we wondered whether other variants in genes encoding EMC subunits might also contribute to early human disease. In our review of the literature, we focused on variants that where likely to disrupt protein function (protein truncations, frameshifts, or splicing disruptions) which included EMC9 and EMC10 (Table 1), prompting us to focus on these EMC components (Jin et al. 2017; Shao et al. 2021; Umair et al. 2020). Patient phenotypes included congenital heart disease (CHD), neurodevelopmental delay (NDD), craniofacial dysmorphology (CFD), umbilical/inguinal hernias, renal anomalies, and limb abnormalities. CFD was the most common phenotype across reported patients with EMC gene variants, though dysmorphologies appeared heterogeneous and varied even amongst patients with the same genotype. Given the prevalence of CFDs in these patients with variants in EMC9/10, we sought to investigate the neural crest given its importance in craniofacial development.

**Table 1:** Mutations in EMC9 and EMC10 in 18 individuals from ten families with congenital anomalies CFD, craniofacial defects; CHD, congenital heart disease; gnomAD Genome Aggregation Database http://gnomad.broadinstitute.org; Het, heterozygous; Hom, homozygous; MAF, minor allele frequency N/A, not applicable; NDD, neurodevelopmental delay

| Subunit Gene | Coding Sequence Change | Amino Acid Change | Effect | gnomAD MAF | Source | Patients/Families | Zygoty | Phenotypes |
| --- | --- | --- | --- | --- | --- | --- | --- | --- |
| EMC9 | c.307C>T | p.Arg103Ter | Stopgain | 1.59E-05 | Jin et al. 2017 | 1/1 | Het | CHD, CFD, NDD |
| EMC10 | c.736delG | p.Gly246fs | Frameshift | 0 | Jin et al. 2017 | 1/1 | Het | CHD, CFD |
| EMC10 | c.679-1G>A | N/A | Splice Acceptor | 0 | Umair et al. 2020 | 2/1 | Hom | CFD, NDD |
| EMC10 | c.287delG | p.Gly96fs | Frameshift | 2.54E-05 | Shao et al. 2021 | 14/7 | Hom | CHD, CFD, NDD |

After neurulation, NCCs delaminate from the neural plate border and undergo an extensive pattern of migration in response to environmental cues. Once they arrive at their destinations, they can differentiate into an extraordinary array of cell types including chondrocytes that establish craniofacial structure. To assess NCC development, we employed whole mount in situ hybridization (WISH) to evaluate sox10 as a NCC marker that defines NCC specification and migration. For these experiments, we exploited an advantage of Xenopus. Due to holoblastic cleavages, injection of one cell at the two-cell stage (Figure 1A) can lead to targeting of either the left or right side of the embryo, providing the unaffected side as an internal control that can be easily identified by co-injecting fluorescent tracers. Thus, we depleted emc subunit genes unilaterally via CRISPR/Cas9 injection in developing embryos. We then examined sox10 expression (Figure 1B). Indeed, sox10 was reduced in affected tissue when emc9 or emc10 was depleted (Figure 1C).

**Figure 1:**
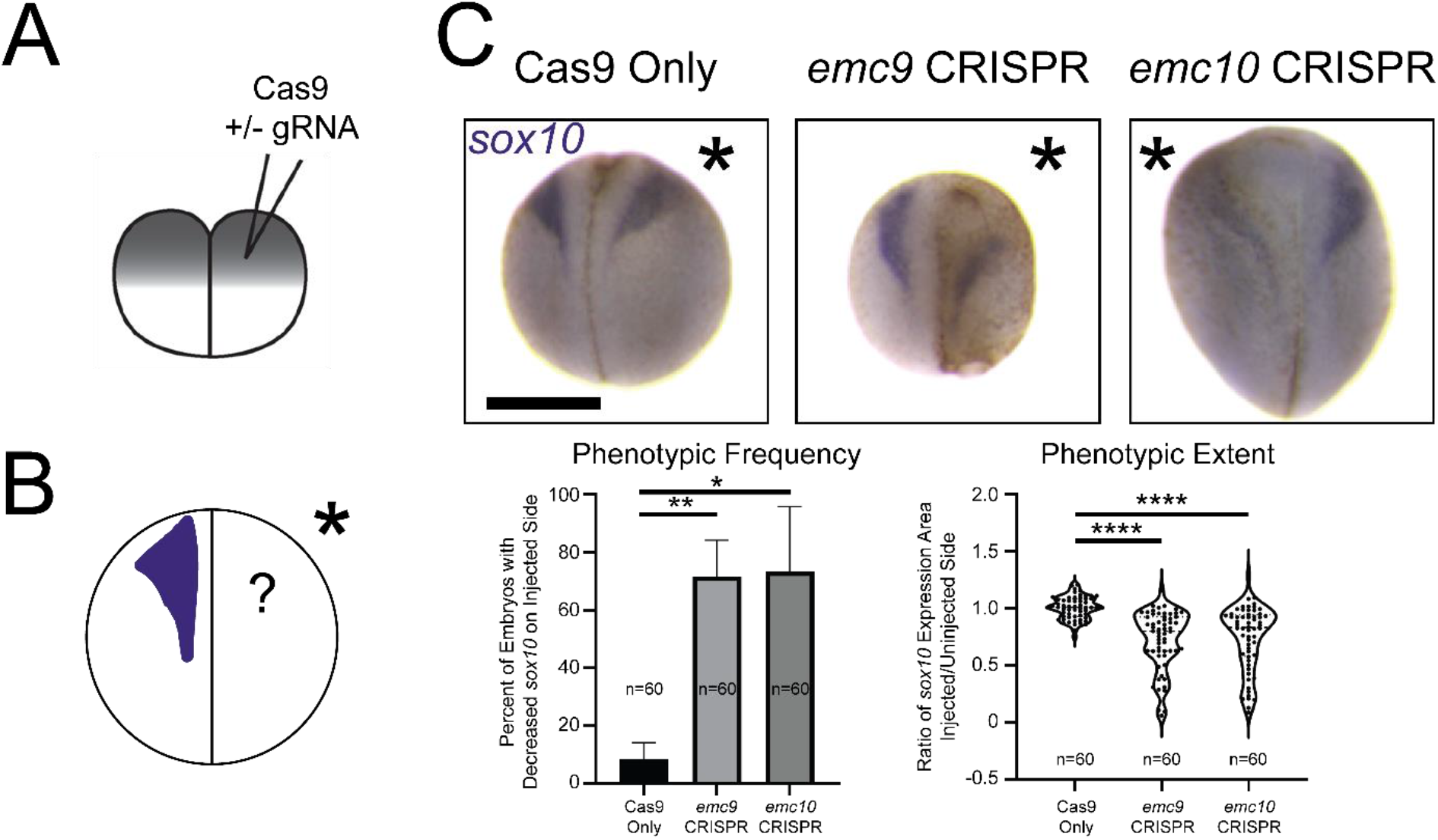
Depletion of emc9 or emc10 in Xenopus results in defective Neural Crest Development. (A) Schematic of injection after holoblastic cleavage at the 2-cell stage (B) Schematic of experimental set up to assess sox10 expression via WISH (C) Representative images and quantitation of WISH for sox10 demonstrates decreased expression at stage 20 in tissues depleted of emc9 or emc10 (n = 60 per condition over 3 replicates; injected halves of embryos indicated by asterisks). Scale bar: 500 μm. Statistical tests carried out as unpaired t-tests for frequency of phenotype and two-tailed t-tests for expression area ratios. *p<0.05, **p<0.01 and ****p<0.0001. Bars indicate mean and SD of replicate values for frequency individual values for area of expression.

We further assessed subsequent development of craniofacial cartilage. To this end we used CRISPR mediated F0 knockout of emc9 or emc10 in Xenopus embryos and assessed craniofacial morphology in injected embryos. As opposed to our assessment of neural crest via sox10, here we employed one cell CRISPR injected embryos to more easily evaluate the overall appearance of craniofacial morphology (Figure 2A). We noted a dysmorphology in craniofacial cartilages for both emc9 and emc10 depletion (Figure 2B). From these studies, we conclude that NCC and craniofacial cartilage development are altered in emc9 and emc10 depleted embryonic tissue.

**Figure 2:**
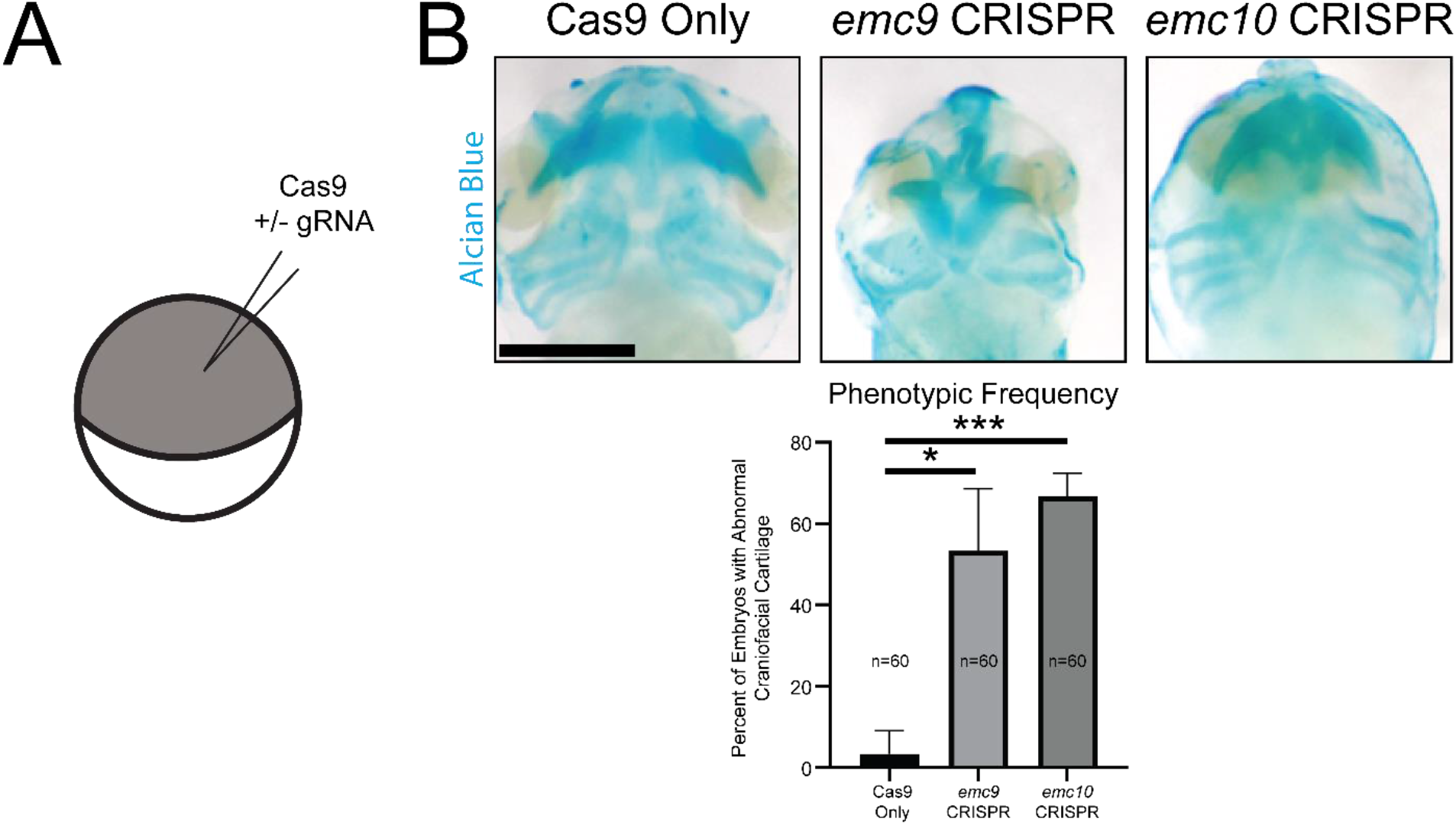
Depletion of emc9 or emc10 in Xenopus results in Craniofacial Cartilage Abnormalities. (A) Schematic of injection at the 1-cell stage (B) Representative images and quantitation of Alcian Blue staining of craniofacial cartilage demonstrates abnormalities at stage 45 in tissues depleted of emc9 or emc10 (n = 60 per condition over 3 replicates). Scale bar: 250 μm. Statistical tests carried out as unpaired t-tests. *p<0.05 and ***p<0.001. Bars indicate SD of replicate values.

As the transmembrane nicotinic acetylcholine receptor (nAChR) has been shown to rely on EMC mediated insertion (Richard et al. 2013) and is crucial in neuromuscular signaling, we next assayed movement in tadpoles depleted of emc9 or emc10. We measured the distance tadpoles moved after stimulation. emc9 or emc10 deficient embryos moved considerably shorter distances than their stage-matched siblings in the control group (Figure 3A). To test whether this might indeed be due to nAChR abnormalities, we performed immunofluorescence for nAChR in tadpole tails, examining the locations where muscle contraction impulses were generated (Figure 3B). In control embryos, the nAChR signal was a sharp arc across the somitic muscle, whereas in emc9 or emc10 depleted embryos, the nAChR signal appeared weaker and more discontiguous. From these results, we conclude that properly localized nAChR is reduced in tadpoles depleted of EMC subunits.

**Figure 3:**
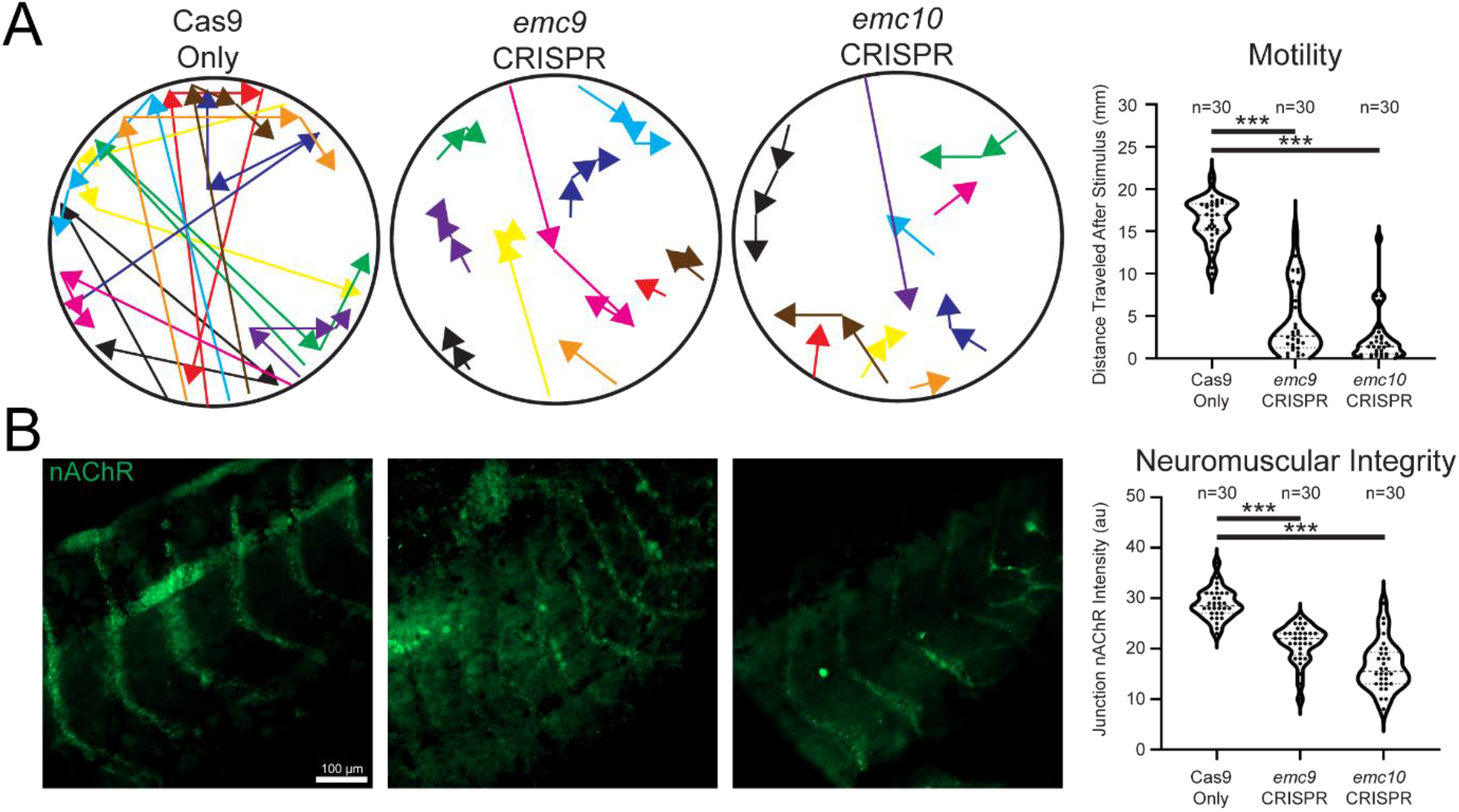
Depletion of emc9 or emc10 in Xenopus affects embryo motility and neuromuscular acetylcholine receptor patterning. (A) Sample traces and measurement of control (n = 30), emc9 depleted (n = 30), and emc10 depleted tadpole movement over 10 seconds after stimulation (different colors differentiate distinct tadpoles) over 3 replicates. (B) Labeling of tail neuromuscular acetylcholine receptor distribution reveals sparse signals in (n = 30) emc9 and (n = 30) emc10 depleted junctions.

As we had previously observed a disruption in WNT signaling in emc1 dysfunction phenotypes, we assayed the levels of transmembrane WNT components. With both emc9 and emc10 depletion we observed a decrease in Fzd7 levels consistent with disruption of Fzd folding (Figure 4A). While this was suggestive of WNT dysfunction on the transmembrane protein level, we sought to assess the downstream effect on WNT signaling that involves nuclear import of β-catenin to enact transcriptional changes. Levels of nuclear β-catenin were decreased in embryos depleted of emc9 and emc10 (Figure 4B).

**Figure 4:**
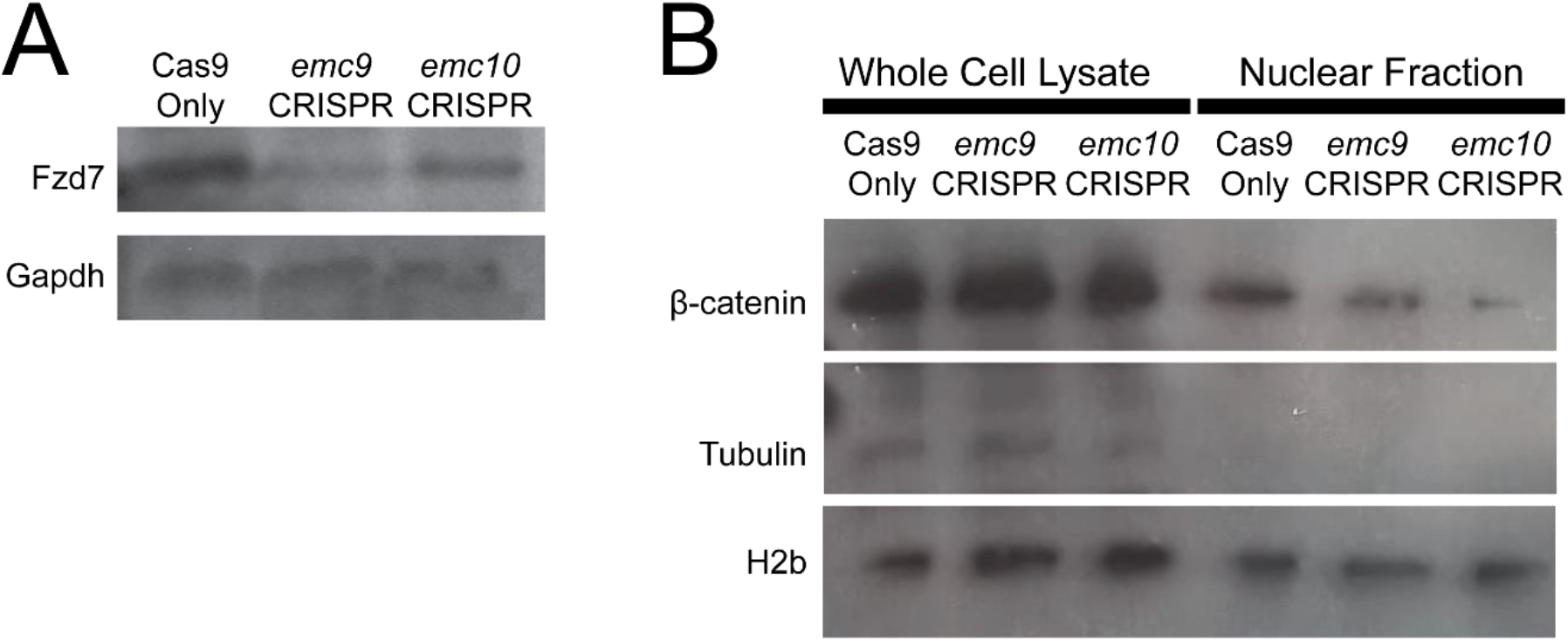
Depletion of emc9 or emc10 in Xenopus affects WNT signaling via depletion of transmembrane protein levels. (A) Immunoblotting for Fzd7 showed a decrease in Fzd7 in pooled (n = 30 per stage per condition) emc9 depleted and emc10 depleted tadpoles as compared with control tadpoles at stage 45. (B) Immunoblotting for β-catenin showed a marked decrease in nuclear β-catenin in pooled (n = 30 per stage per condition) emc9 depleted and emc10 depleted tadpoles as compared with control tadpoles at stage 45.

## Discussion

As the EMC is both evolutionarily conserved and utilized in all known cell types, we believed that dysfunction in different subunits would likely result in similar phenotypes via closely related mechanisms. Yet, recent work assessing multiple subunits of another large complex, the nuclear pore complex, uncovered distinct phenotypes and mechanisms leading to congenital disease (Braun et al. 2018; Braun et al. 2016; Chen et al. 2019; Del Viso et al. 2016; Marquez et al. 2021; Miyake et al. 2015; Muir et al. 2020). Thus, modeling loss-of-function phenotypes of subunits in a large complex has proven a fruitful endeavor.

The observed phenotypes in patients with likely damaging variants of EMC9 and EMC10 closely resembled those of patients with EMC1 variants. Yet, the inheritance pattern and phenotype correlation among variants in EMC9 and EMC10 appears distinct. Heterozygous loss-of-function variants in EMC1 resulted in CHD while homozygous putative loss-of-function variants resulted in a syndromic NDD phenotype. The heterozygous putative loss-of-function variant in EMC9 may contribute to both CHD and NDD phenotypes while both heterozygous and homozygous putative loss-of-function variants in EMC10 appear to contribute to either CHD or NDD or both (Table 1). Larger clinical cohorts of patients with EMC variants will be essential to fully decipher these phenotype-genotype correlations.

Given these observations, we concluded that modeling loss-of-function in EMC9 and EMC10 in Xenopus would be useful in further determining the pathogenicity of loss-of-function in these genes. By assessing NCC development in emc9 and emc10 depleted tissues through the late NCC marker sox10, we observed mis-patterning which indicates developmental dysfunction within the NCC lineage. Later assessing craniofacial cartilage, we uncovered dysmorphology indicative of impaired craniofacial cartilage establishment downstream of impacts on NCC development. Our investigation of neurological dysfunction further supported the role of EMC subunit dependence for proper neuromuscular signaling. It therefore appears that emc9 and emc10 loss-of-function affects the same tissues as emc1 loss of function, including NCC derivatives. Indeed, given the overlap of the observed phenotype in our now multiple EMC subunit loss-of-function gene models, it appeared likely that dysfunction leads to congenital disease via a similar deficiency in protein folding.

Visualization of protein localization of nAChR and decreases in Fzd7 further support the role of transmembrane protein misfolding as the basis for EMC subunit loss of function resulting in phenotypes observed in our Xenopus model and patients. In particular, it appears that the downstream diminution of WNT signaling likely plays a crucial role in these phenotypes. We observed both loss of transmembrane components of WNT signaling as well as evidence of disrupted downstream WNT signaling.

Our understanding of the EMC has flourished amidst recent studies on the structure, molecular function, and determination of the client proteins of this crucial complex (Chitwood et al. 2018; Guna et al. 2018; Miller-Vedam et al. 2020; O’Donnell et al. 2020; Pleiner et al. 2020; Shurtleff et al. 2018; Tian et al. 2019). While the phenotypes observed in our Xenopus emc9 and emc10 loss-of-function models resembled those of emc1 loss of function this is somewhat surprising given the relative importance of EMC subunits. Studies of EMC function have identified similar structures and possibly even a similar evolutionary origin for EMC8 and EMC9 (Bai et al. 2020; O’Donnell et al. 2020; Pleiner et al. 2020; Tian et al. 2019; Wideman 2015). Given this potential redundancy between EMC8 and EMC9 we expected a potentially less severe phenotype in our emc9 loss-of-function model. As this was not apparent in our studies, it may point to a distinct role for EMC9 during the window of development that we interrogated.

As additional variants in EMC subunit genes are identified, we may begin to better understand a critical question about the role these proteins play in essential aspects of cell biology. Thus far, only putatively damaging variants in EMC1, EMC9, and EMC10 have been observed in patients. These genes encode portions of the EMC that are not considered “core” components. EMC2, EMC5, and EMC6, the core components of the EMC are required not only for proper folding of client EMC proteins but also for the establishment of the EMC itself (Volkmar et al. 2019). As no disease-causing variants have been observed in the genes encoding these subunits, perhaps this indicates that we have observed EMC1, EMC9, and EMC10 variants due to a greater tolerance for mutation albeit causing severe disease.

In summary, our results indicate that disruption of additional subunits of the EMC complex support a model of human disease via dysfunction in the neural crest and derived tissues stemming from transmembrane protein misfolding. This conclusion is readily evident in our Xenopus model and extends the implications of our previous work on the pathogenesis of dysfunctional EMC disease. Indeed, EMC subunit related diseases may be best understood as a category of developmental foldopathies. Future therapeutic approaches could aim to alleviate the excess of misfolded proteins as a strategy to ameliorate disease burden, given the difficulty of targeting WNT signaling itself. These mechanistic insights may serve to improve treatment and optimize care for future patients with EMC variants.

## Author Contributions

JM and MKK conceived, designed, and analyzed all experiments. JM performed all experiments. The manuscript was written by JM and MKK.

## Acknowledgements

We thank Michael Slocum for animal husbandry. JM was supported by the Yale MSTP NIH T32GM07205 Training Grant, the Yale Predoctoral Program in Cellular and Molecular Biology T32GM007223 Training Grant, and the Paul and Daisy Soros Fellowship for New Americans. This work was supported by the NIH R01HD081379 to MKK.

## Notes

### Competing Interest Statement

MKK is a founder of Victory Genomics, Inc

