## Supplemental Table 1 for "Expanding EMC Foldopathies: Topogenesis Deficits Alter the Neural Crest"

| Reagent | Vendor | Catalog | Dilution | Species |
|---|---|---|---|---|
| α-BUNGAROTOXIN (labeling of NAChRs) AF488 | Thermo Fisher Scientific | B13422 | 1:1000 | *Bungarus multicinctus* |
| FZD7 | Abcam | ab64636 | 1:1000 | Rabbit |
| β-Catenin | Santa Cruz | sc-7963 | 1:1000 | Mouse |
| γ-Tubulin | Sigma Aldrich | T6557 | 1:1000 | Mouse |
| H2B | Abcam | ab1790 | 1:1000 | Rabbit |

Supplemental Table 1: Reagents and antibodies used for immunohistochemistry and immunoblotting
